# Thermal selection shifts genetic diversity and performance in blue mussel juveniles

**DOI:** 10.1101/2024.12.12.628131

**Authors:** Jennifer C. Nascimento-Schulze, Jahangir Vajedsamiei, Tim P Bean, Lisa Frankholz, Reid S. Brennan, Frank Melzner, Robert P Ellis

## Abstract

Mussels from the genus Mytilus, key inhabitants of the benthos, are important for the aquaculture industry and one of the most sustainable sources of animal protein available. Species within the *Mytilus edulis* complex (*M. edulis, M. galloprovincialis* and *M. trossulus*) are commonly found in temperate regions globally and can easily hybridise whenever their geographic distributions overlap. In the Baltic Sea, populations are formed by *M. edulis* and *M. trossulus* hybrids with low levels of *M. galloprovincialis* introgression. Given the economic and ecological relevance of mussels, this study aimed to investigate mechanisms through which their resilience towards global warming may be fast-tracked. For this, we developed two cohorts of juvenile mussels (i.e. recently settled animals) from the Baltic Sea (Kiel, Germany), one exposed to an extreme heat event early in life and one naïve to this stressor. Both cohorts were then exposed to experimental temperatures at the proposed upper thermal limit for this population, 21^°^C to 26^°^C, with animal performance measured after 25 days. We then assessed the impacts of thermal stress on the genetic composition of each cohort by genotyping 50 individuals using the blue mussel 60K SNP-array. We observed a significant increase in *M. edulis* genotypes together with a decrease in *M. trossulus* in the S cohort in comparison with NS juveniles. We also found that exposure to high temperature has an effect on the performance of mussel cohorts, reducing dry tissue weight of the selected individuals. Results from this study provide relevant insights on how selection through thermal stress impacts performance and genetic composition of blue mussel juveniles, with key implications for understanding and managing mussel populations under future warming scenarios.

## Introduction

Ongoing climate change is shifting abiotic and biotic patterns in the ocean, affecting the fitness of marine organisms and the functioning of marine ecosystems worldwide (IPCC 2022). Sea surface temperatures (SST) in the Baltic Sea have risen more rapidly than in other marginal Seas (Belkin, 2009). Over the past decades, the region has experienced an increase in SST of more than 0.3°C, with warming trends being more pronounced during summer months compared winter (Meier et al. 2022). Heatwave events, which are increasing in duration and intensity due to climate change, can further exacerbate warming, leading to SST extremes (Frölicher et al., 2018). For instance, in 2018, the warmest summer since 1990 (Naumann et al., 2018), heatwaves raised temperatures by more than 4°C above the 28-year mean (1990-2018) in the southern portion of the basin (Meier et al., 2022). Shifting temperature regimes, either through modified mean seasonal temperatures or isolated shorter extreme events, can contribute to community shifts and set off a chain reaction of biodiversity loss, pushing this ecosystem towards an alternate state (Johannesson et al., 2011; Pansch et al., 2018; Reusch et al., 2005).

Blue mussels are a foundation species of the benthos, providing ecosystem services such as nutrient cycling and increased habitat complexity through enhanced spatial structure for associated species (Johannesson et al., 2011; van der Schatte Olivier et al., 2020), as well as being an important aquaculture and fisheries resource (Avdelas et al., 2021). The three species in the blue mussel *Mytilus* complex, *M. galloprovincialis, M. edulis* and *M. trossulus* hybridise in areas of overlapping geographical distribution (Fraïsse et al., 2016). In the Baltic sea, blue mussel populations are a highly introgressed swarm composed of all the three species (Vendrami et al., 2020). These populations already cope with environmental pressures, such as warming, at levels that most coastal areas are not expected to experience until the end of the century (Reusch et al., 2018). However, little is known concerning the implications of thermal stress in these populations on a genomic level.

Temperature is a major factor regulating the physiology of ectotherms (Somero et al., 2017). In these organisms, the relationship between physiological performance and body temperature can be described by an asymmetric thermal performance function (Angilletta, 2009). When environmental temperatures fall within the tolerance range of a species/population, because of positive thermodynamics on enzymatic activity, metabolic rates rise, and an organism increases its respiration rate and feeding rates to supply the enhanced energy demand. However, when temperatures exceed critical thermal maximum (CT_max_) of a species/population, a set of responses at the cellular level limit survival (Pörtner and Farrell, 2008; Somero, 2010). During acute heat exposure, mussels metabolically adjust to elevated temperatures by increasing their metabolism, reducing high energy demanding processes (e.g. growth) and/or supplementing aerobic with anaerobic metabolism to cover rising metabolic demands (Anestis et al., 2007a; Braby and Somero, 2006; Tagliarolo and McQuaid, 2015; Vajedsamiei et al., 2021b; Zittier et al., 2015). These strategies can guarantee short-term survival in non-optimal thermal conditions. Long-term exposure to intermediate warming (24°C) leads to metabolic suppression, through the lowering of pyruvate kinase (PK) glycolytic enzyme activity in *M. galloprovincilis*. In addition, *M. galloprovincialis* can respond to sub-lethal warming with increasing heat-shock protein transcription when temperatures rise above ca. 22°C (Feidantsis et al., 2020). Long-term exposures to extreme temperatures of 26°C and above trigger a reactivation of anaerobic metabolism (Anestis et al., 2007) to fuel energy intensive heat-shock protein production (Hawkins, 1985), and other cellular rescue processes (Feidantsis et al. 2020). However, survival at temperatures exceeding an individual’s CT_max_ is time-limited. The high energetic demands trigger a temperature-induced mismatch between metabolic supply and demand, hindering performance (Pörtner, 2012; Ritchie, 2018) and leading to mortality (Anestis et al., 2007, Feidantsis et al. 2020).

The capacity to adjust metabolic rate in response to temperature is a critical survival trait and varies across species (Pörtner, 2012; Vajedsamiei et al., 2021b). For example, the upper thermal limits for maintaining a positive Scope for Growth (i.e. the extra energy available for growth after meeting basic survival needs) in *M. trossulus, M. edulis* and *M. galloprovincialis* acclimated to summer temperatures from the north western Pacific and north eastern Atlantic coasts, are ca. 17°C, 23°C and 30°C, respectively (Fly and Hilbish, 2013). These results indicate that at elevated temperatures, *M. trossulus* genotypes have a lower performance in comparison to the other two species, with likely implications for growth, survival and reproductive output. However, it remains unclear if the genetic contributions from different *Mytilus* species within hybrid populations can affect their responses to thermal stress. In this study, we addressed this knowledge gap via a two-step experimental approach (Figure 1). First, to assess the implications of thermal stress on the genetic makeup of mussel populations, we collected spat (i.e. recently settled juveniles) from a single settlement event within Kiel Fjord and developed two distinct cohorts, one selected for thermal tolerance via an acute but intense exposure to a thermal stress event (30°C for 47 h), and one naïve to this stressor. Individuals from both cohorts were genotyped with the 60K Blue mussel SNP-array (Nascimento-Schulze et al., 2023). We hypothesised that temperature selection would create a genetically distinct cohort, favouring genotypes more resilient to higher temperatures. Specifically, we expected the fraction of *M. galloprovincialis* and *M. edulis* alleles to increase following temperature selection. Second, we tested the impacts of post-selection thermal stress on mussel performance. For this, we exposed individuals from both developed cohorts to constant Baltic summer temperature scenarios, ranging from current to forecasted end-of-century summertime extremes, over a 25-day period, and assessed whole organism performance in shell growth and dry tissue gain. We hypothesized that heat selected lines would outperform non-selected lines at higher temperatures.

**Figure 1:**
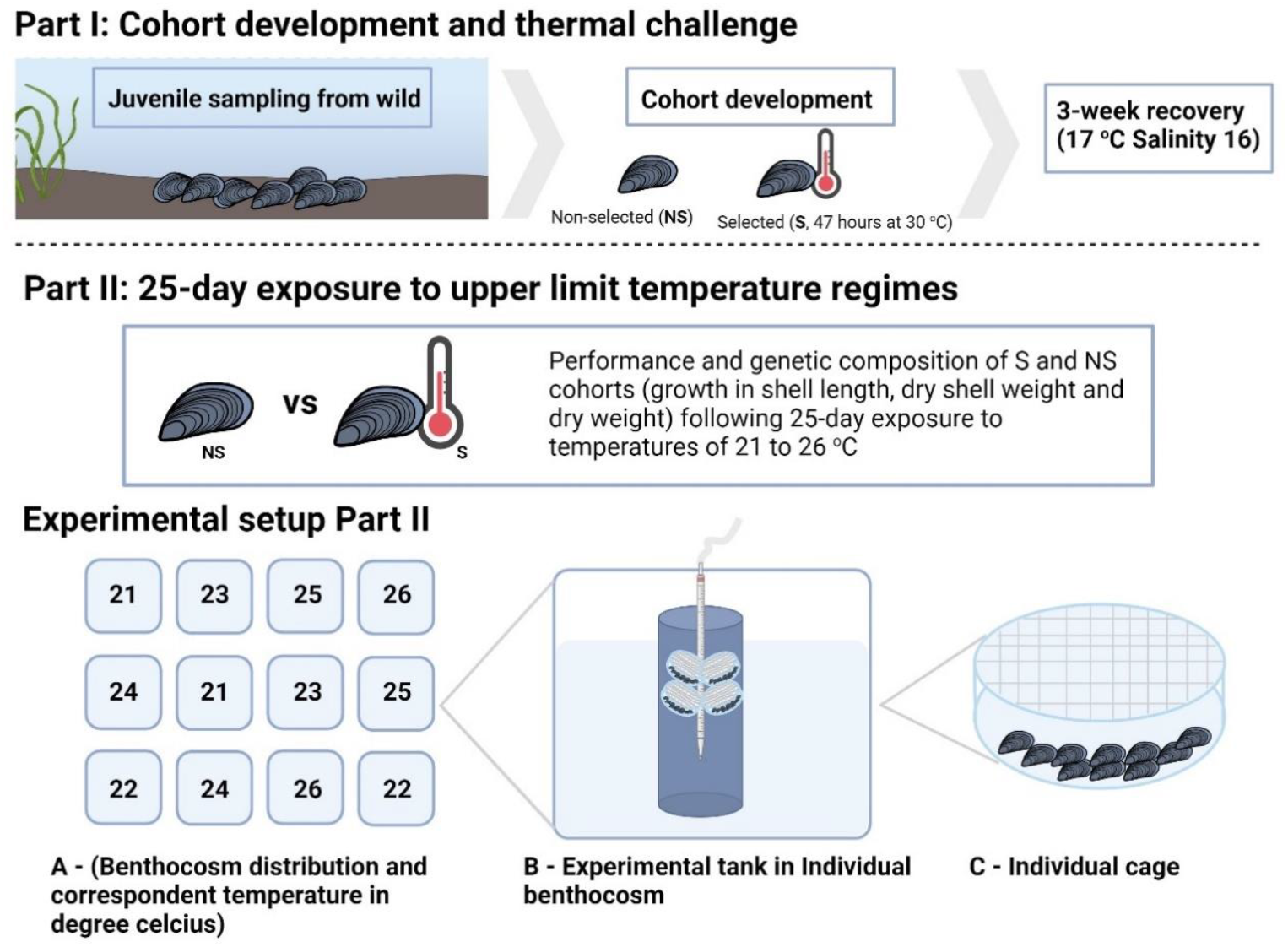
Overview of the experiment specifying the main steps. Part I describes the development of both cohorts (selected and non-selected) following selection after a simulated heatwave event. Part II outlines the experimental set-up for the 25-day thermal challenge conducted in a controlled indoor benthocosm system, with a representation of the benthocosm layout (A), a schematic representation of the tank setup within individual benthocosms; experimental cylinders filled with seawater, aerated via a plastic pipette, and holding four chambers containing mussels (B); and finally an individual mussel chamber (C).

**Figure 2:**
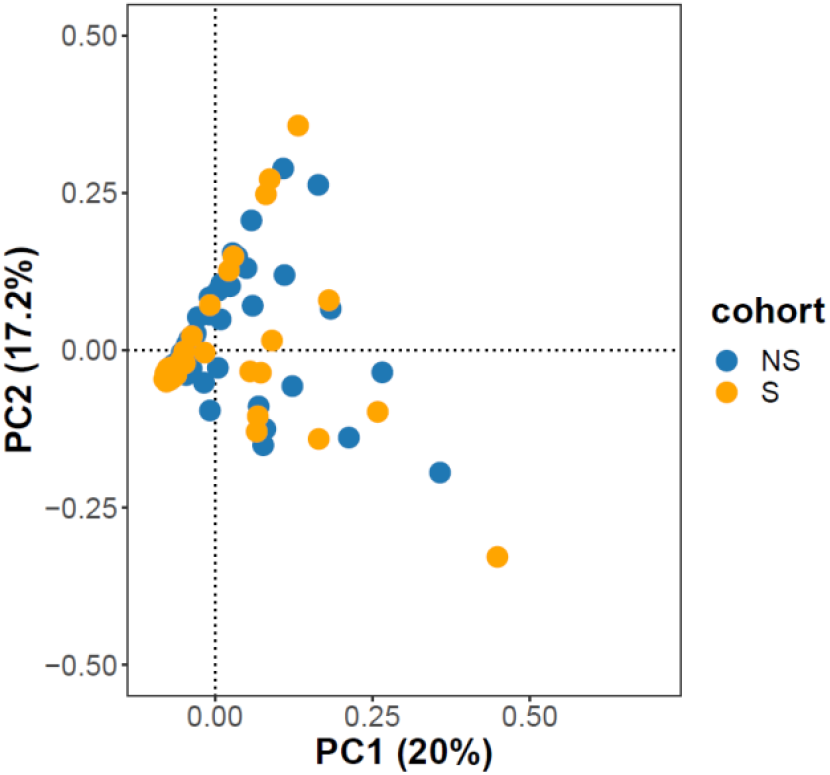
Scatter plots of individual variation in PC 1 and 2 scores resulting from PCA applied to the blue mussel juveniles from Kiel fjord, Germany. Selected (S) and Non-selected (NS) individuals were genotyped with the blue mussel multi-species 60K SNP array. The proportion of overall variation explained by each PC are given in percentages.

## Materials and Methods

### Development of selected and non-selected juvenile cohorts

#### Selection event

Approximately 500,000 mussel spat (i.e. settled individuals) were collected by hand from subtidal Kiel Fjord (54.328588, 10.148210, Baltic Sea, Germany) on July 28^th^, 2021 from approximately 0.5 m depth, and gently sorted using a 1-mm mesh size plastic sieve suspended in a 10 L bucket of seawater. On the following day, around fifty thousand spat were evenly distributed into five separate 10 L tanks (∼10,000 spat per tank). These naïve spat were kept at 17°C (temperature-controlled room at GEOMAR) with 16 PSU natural seawater filtered through a 0.5 μm mesh. For the next eight days, husbandry practices involved 50% water exchanges daily and the addition of 100 mL of *Rhodomonas salina* (∼1-2 x 10^6^ cells mL^-1^) as food. The remaining 450,000 spat were then thermally challenged: first, being evenly distributed among 16 x 2 L tanks (ca. 25,000-31,000 per tank) filled with 600 mL of 16 PSU natural seawater, filtered with a 0.5 μm mesh, and exposed for 30 minutes to a non-lethal elevated temperature (26.5°C), with constant aeration supplied through 5 mL plastic pipette tips. The experimental tanks were placed into four steel water baths that maintained target temperatures with a deviation of less than 0.1°C (model Haake SWB25, Thermo Scientific). A further 1 L of filtered sea water (FSW; filtered across a series of filters 10- and 1-micron mesh) at 30°C was added to each tank, water baths were set to 30°C and target temperatures were reached within ∼15 min. Spat were then kept at this temperature for 47 hours.

Three days post collection, all heat-treated spat (dead or alive) were washed in FSW to remove tissue of dead mussels and then transferred from the 16 x 2 L tanks into 4 x 10 L tanks with FSW (salinity 16 PSU, temperature 17°C ±0.2°C, constant aeration), located adjacent to the tanks housing the naïve animals in a temperature-controlled room. For the first two days post transfer, water changes (∼95% of volume) were undertaken daily to remove the organic remains of dead animals followed by the addition of 200 mL of *R. salina* (∼1-2 x 10^6^ cells mL^-1^) to each tank. Over the following nine days, husbandry practices shifted to daily 50% water exchange and the addition of 100 mL of *R. salina* (∼1-2 x 10^6^ cells mL^-1^). Heat-selected spat were then stained with calcein green and naïve animals with calcein blue (50 mg L^-1^ on day 1, and 25 mg L^-1^ on day 2-9), added immediately after the water changes, to fluorescently label live individuals. Both chemicals are non-toxic to mussels.

At 14-days post collection, heat-selected spat from the four tanks were mixed and three 500 μL sub-samples of mussels were pipetted into a 0.5 mL Eppendorf tube, with survival rates assessed. Due to the calcein shell labelling procedure, living spat appeared bright green under the stereomicroscope equipped with standard GFP-filter sets (Leica M165 FC), and 600 living spat were found and kept for the subsequent steps. In addition, approximately 600 spat were randomly sorted from the ∼ 50,000 naive spat kept in the five maintenance tanks at constant temperatures of 17°C. No mortalities were observed in the naive group. Each group of 600 naïve or selected spat was then equally distributed among four 50 mL containers (150 individuals in each container, 4 containers per spat line), which were closed with mesh (300 μm). Two of these containers, one containing selected and one containing the non-selected spat were kept in one 10 L aquarium (‘conditioning tank’, total of four tanks) for seven days at 17°C and 16 PSU, fed every day with an initial concentration of ca. 8,000 *R. salina* cells mL^-1^ to ensure comparable condition between spat from the two different treatment groups for the following experiments.

In the following sections of this manuscript, spat surviving the thermal stress exposure are referred to as ‘selected’ (S), whilst spat not exposed to thermal stress are termed ‘non-selected’ (NS). These spat were exposed to a thermal challenge in the Kiel Indoor Benthocosm facilities (KIB, Pansch and Hiebenthal, 2019).

#### Second stress exposure

Following the initial development of S and NS spat cohorts, 960 individuals (480 S and 480 NS) were randomly selected for a follow-up experiment, in which we assessed the impact of thermal stress on juvenile performance. Beginning on August 20^th^, spat were exposed to 5 different temperature treatments (21, 22, 23, 24, 25 and 26 °C) over a 25-day period. The 26°C treatment represented a predicted end-of-century peak daily marine heatwave temperature in the study region , sub-lethal to mussels for periods of <1 month . The 21°C treatment represents a present-day maximum summer temperature, which is in the optimal range for mussel growth. The intermediate temperatures within the experimental range were selected to identify any subtle, but significant, alterations in thermal performance or thermal breakpoints in larval mussel performance in this study. The experiment was conducted using the KIB. In this facility, 12 x 600 L polyethylene tanks (i.e. benthocosms) equipped with GHL Profilux computers and thermal sensors (GHL GmbH), controlling heaters and chillers (Aquamedic), enabled the maintenance and recording of water temperature. Two experimental cylinders were assigned to each of six experimental temperatures. KIB tanks were used as water baths and housed the12 animal incubation cylinders (two per water bath). Animals were placed into plastic cages (30 mm O.D, approximately 10 mm length) sealed on the top and the bottom with 300 μm nylon mesh (Figure 1). Experimental cages were placed into 14 L plastic cylindrical tanks, allowing water flow and food uptake to occur. Twenty juveniles were kept in each cage, with each treatment (S and NS) having two replicate cages per experimental tank. Thus, a total of 80 S and 80 NS animals were incubated at each temperature. Cages were tied to 15 mL pipettes that were used for aeration at the same water column height, providing similar environmental conditions between replicates of each treatment (Figure 1).

Initial shell length was measured in a sub-sample of 60 S and 60 NS individuals, which were removed from the conditioning tank before being distributed into experimental cages on day 0, one day prior to the start of the 25-day thermal exposure experiment. The impact of elevated temperature on mussel spat was assessed by measuring larval performance via growth as shell length, dry tissue weight and dry shell weight on day 25, at the end of the challenge, from at least 10 animals per cage. Raw data values in the results section are presented as mean ± standard deviation.

#### Husbandry

Throughout the duration of this experiment, spat were fed with monocultures of *Rhodomonas salina*, added at a concentration of ∼ 8,000 cells mL^-1^ (Riisgård et al., 2011) to the experimental cylinders after every water exchange. Cell concentration was measured daily (particle size 5-8 um) using a Coulter counter (Coulter Z2, Beckman Coulter GmbH), in all experimental tanks. If *R. salina* concentration in any of the experimental tanks dropped to threshold values in the range of 1,000 cells ml^-1^, full water exchanges were performed in all the tanks in order to ensure that *R. salina* was constantly available in the water column at optimal concentrations. Water changes were conducted with FSW every 3^rd^ day (days 1-7), every other day (days 8-20), and daily after day 21 (Suppl. Table 1).

#### Growth, dry body mass and shell mass assessment

Shell length was measured using vernier callipers. Dry tissue weight and shell weight were measured as previously described. Briefly, sampled spat were killed in a microwave (400 watts, 30 s), and their soft tissue was removed from their shell under a stereomicroscope. Shell and soft tissues were separately placed individually into pre-weighed tin foil boats dried at 80°C, for a minimum of 72h, and reweighed to determine the dry tissue weight and shell weight (mg) by subtracting the value of the empty foil boat from the foil boat containing shell or soft tissue. Raw data values in the results section are presented as mean ± standard deviation.

### Assessing structure and genomic relatedness of the two cohorts

The genetic composition of S and NS cohorts was analysed using 50 S and 50 NS spat collected on day 0 of the second experiment, i.e. the day prior to the start of the 25-day exposure challenge, with sampled spat preserved in 90% ethanol. Samples were sent to Identigen Ltd for DNA extraction and sequencing, using the 60K blue mussel SNP-array (Nascimento-Schulze et al. 2023).

To analyse the resulting sequencing data, a quality control filter for marker call rate (CR) of > 95% and sample CR >90% was applied using PLINK v1.9. We applied a minor allele frequency (MAF) filter < 0.01. A principal component analysis (PCA) was applied to genotypes generated with the array, pruning out putatively linked loci using a window size of 50 Kb, a step size of 10 Kb and an r^2^ threshold of 0.1 with PLINK v1.7. We explored introgression in spat by applying an admixture analysis (admixture v1.3). In this analysis, the best fitting number of clusters representing the ancestry of a population is estimated where *k* has the lowest cross-validation error value. For this study, we tested *k* values between 2 and 10. In order to allocate a species to the ancestry cluster generated in this first admixture analysis, we ran an additional admixture analysis including 5 additional populations, Kiel (Baltic Germany, GK), Ahrenshoop (Baltic Germany, GA), Finland (Baltic Finland, FIN), Bude (North Devon coast of the UK) and Budleigh (South Devon coast of the UK). These populations have been genotyped with the blue mussel multi-species 60K array in previous studies (Nascimento-Schulze et al. 2023; Nascimento-Schulze et al unpublished data), and their ancestry is known to be *M. edulis/M. trossulus* hybrids for GK and GA and FIN populations, with an increased frequency of *M. trossulus* genotypes in FIN, *M. galloprovincialis* ancestry dominating Bude and *M. edulis* prevails in Budleigh. Admixture results of this data set are presented in Suppl. Table 2 and visualised in Suppl. Figures 3 and 4.

### Statistical analysis

#### Spat performance

Shell length data collected from individuals at day-0, prior to the start of the 25-day thermal challenge, was first tested for normality, and homogeneity of variance, using Shapiro-Wilk and Levene’s tests respectively, before a one-way ANOVA analysis was applied to test for the effects of temperature selection.

We investigated the effects of temperature on the dry tissue weight, shell weight and shell length of S and NS individuals following the 25-day thermal challenge. These effects were modelled as generalised additive mixed models (GAMMs) using *gam* function from *mgcv* package. In the GAMMs, *temperature* was defined as a smooth-effect predictor (with linear and/or nonlinear effects) and *selection* as an ordered factor. Then, using t-statistics and f-statistics (Wald test), the intercept (or the mean) and the slope or nonlinearity (effective degrees of freedom, *edf*) of the reference level smoother (*non-selected*) were each compared to zero, respectively; and, the treatment level (*selected*) smoother’s estimates were then compared to those of the reference level. In addition, the random effects of *water bath* and *cage*, as possible causes of residual dependence, were included, and for unbiased estimation of variance components, GAMMs were fitted using restricted maximum likelihood.

The models were first fitted with the *identity* link function with an assumption of residuals having a gaussian distribution around the predicted means. The assumptions regarding the distributions of residuals were checked via the *DHARMa* package. As it was determined that the residual assumptions were violated, we fitted the models assuming the scaled t-distribution (*scat*) of residuals and found this assumption to be valid. For shell length, the *log*_*10*_ link function was also needed to validate the residual assumption. In terms of the Akaike Information Criterion (AIC), the *scat* models were also superior to the gaussian models. We used the *predict* function from *car* package to predict the means and confidence intervals of responses to predictors. A p value of < 0.05 was assumed as the significance level for all analyses. All analyses were conducted in R v4.3.1. Raw data values in the results section are presented as mean ± standard deviation.

#### Population genetic composition

We compared the genetic composition of the two mussel cohorts (S and NS) to investigate potential differences in ancestry proportions resulting from the selection event. For this, we specified a Bayesian regression model assuming a Logistic-Normal distribution for the ancestry proportions, utilizing the *brms* package (Bürkner 2017) in R v4.3.1. The Bayesian Logistic-Normal Model was chosen as it allows for the comparison of mean proportions and variability within the compositional data, while accounting for correlations between components, making it a robust choice for our analysis. The model was fitted using the No-U-Turn Sampler (NUTS) within the Hamiltonian Monte Carlo framework. A thorough description of the models tested and applied is available in the supplementary information.

## Results

### Impacts of increased temperature on cohort survival and genetic makeup

At 10 days post thermal challenge, survival rate of the selected spat cohort was approximately 8%. No death was observed in NS mussels during this period.

From the 15,049 genotyped loci with a CR > 95%, a total of 11,288 loci were retained after applying the MAF > 0.01. After variant pruning, a total of 10,085 putatively non-linked loci were used for PCA and Admixture analysis.

The PCA analysis revealed no obvious stratification between S and NS cohorts (Figure 3). In this analysis, PC1 explained 20% of the variance, whilst PC2 and PC3 explained 17.2% and 7.64%, respectively.

**Figure 3:**
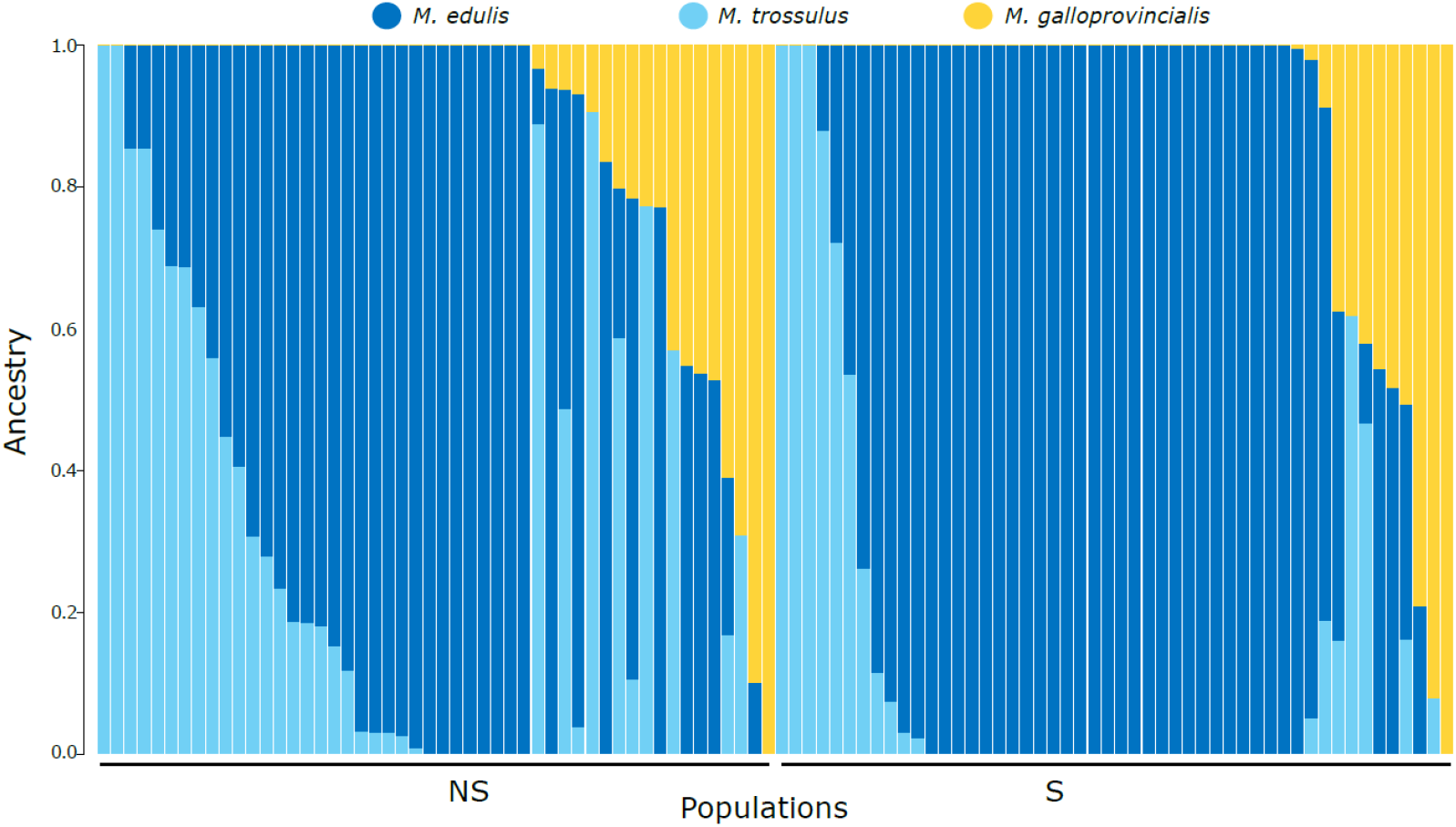
Results of genetic admixture analysis and ancestry inference. Cluster membership coefficients (Q) where each individual is represented by a column partitioned into segments of different colour, the length of which indicate the posterior probability of membership in each cluster. Solid bars represent an individual from a single species background.

The best fitting number of ancestry clusters in the admixture analysis was three (Suppl. Table 2). Results from the admixture coefficients (Q) inferring the three clusters are presented in Figure 3. Based on the analysis, including the control populations (Suppl. Figures 3,4), we could infer the species for each of the three clusters present in the S and NS cohorts, with the dark-blue cluster representing *M. edulis* ancestry, light-blue *M. trossulus* and yellow *M. galloprovincialis*. In this analysis, we observe that most of the genotypes are allocated to *M. edulis*, followed by *M. trossulus* and finally *M. galloprovincialis*.

In our Bayesian analysis, the Logistic-Normal model estimated population-level effects for both q2 (*M. edulis*) and q3 (*M. galloprovincialis*), revealing important trends. For q2, the intercept for the NS was estimated at 2.40 (95% CI: 0.36 to 4.37), while the effect of population S was 3.75 (95% CI: 0.9 to 6.57). This indicates a significant increase in the proportion of *M. edulis* in the S population compared to NS, though the wide credible interval suggests some uncertainty regarding the magnitude of this increase. For q3, the intercept was -3.21 (95% CI: -5.04 to -1.39), and the population S effect was 2.08 (95% CI: -0.56 to 4.74), suggesting a low baseline proportion of *M. galloprovincialis* in the S population. Although there is a positive trend for an increased proportion of *M. galloprovincialis* in population S the credible interval includes negative values, indicating that the evidence for this increase is weak. The family-specific parameters showed significant variability in the proportions of both *M. edulis* (σq2 = 7.24, 95% CI: 6.32 to 8.33) and *M. galloprovincialis* (σq3 = 6.73, 95% CI: 5.87 to 7.74) across individuals, pointing to considerable heterogeneity within the populations. Notably, the model estimated a moderate positive correlation between q2 and q3 (correlation = 0.48, 95% CI: 0.33 to 0.62), suggesting that as the proportion of *M. edulis* increases within a population, there is a tendency for the proportion of *M. galloprovincialis* to increase as well, though this relationship is not particularly strong.

### Impacts of increased temperature on shell length, dry tissue weight and shell mass

No significant difference in shell length was observed between S (2.02 ± 0.33 mm) and NS (1.99 ± 0.32 mm) cohorts at day-0, prior to the start of the 25-day thermal challenge (Suppl. Figure 5, Suppl. Table 3, p > 0.05).

Exposing spat to different experimental temperatures over a 25-day period did not result in a significant difference in shell length between S and NS cohorts (Figure 4B, Table 1, p > 0.05). The mean shell length of S and NS cohorts at day 25 was 2.33 ± 0.52 mm and 2.47 ± 0.67 mm, respectively, suggesting a marginal if non-significant (p = 0.0663) effect of selection on subsequent growth. The smooth effect of temperature was significant overall, as shown by the NS group (p = 0.0006), but was not significantly different between the S and NS groups (p = 0.1260), with NS juveniles at 26°C smaller (2.04 mm ± 0.45) than those at 21^°^C (2.37 ± 0.49 mm). For dry tissue weight analysis, the intercept for the S group, representing its difference from the NS baseline, was significantly lower (p = 0.0001; Table 1), indicating an effect of prior selection on dry tissue mass in all temperature treatments in the follow-up experiment. The smooth effect of temperature was significant overall, as shown by the non-selected group (p = 0.0001), but was not significantly different between the selected and non-selected groups (p = 0.5926). Observed dry tissue weight of NS animals decreased from 0.16 ± 0.06 mg at 21°C to 0.06 ± 0.10 mg at 26°C and dry tissue weight of S juveniles decreased from 0.14 ± 0.11 mg in 21°C to 0.05 ± 0.15 mg in 26°C. Finally, no significant impact of temperature or selection was observed on shell weight of individuals (Figure 4B, Table 1), with shell weight of S spat 0.79 ± 0.5 mg and NS spat of 0.77 ± 0.36 mg.

**Table 1:**
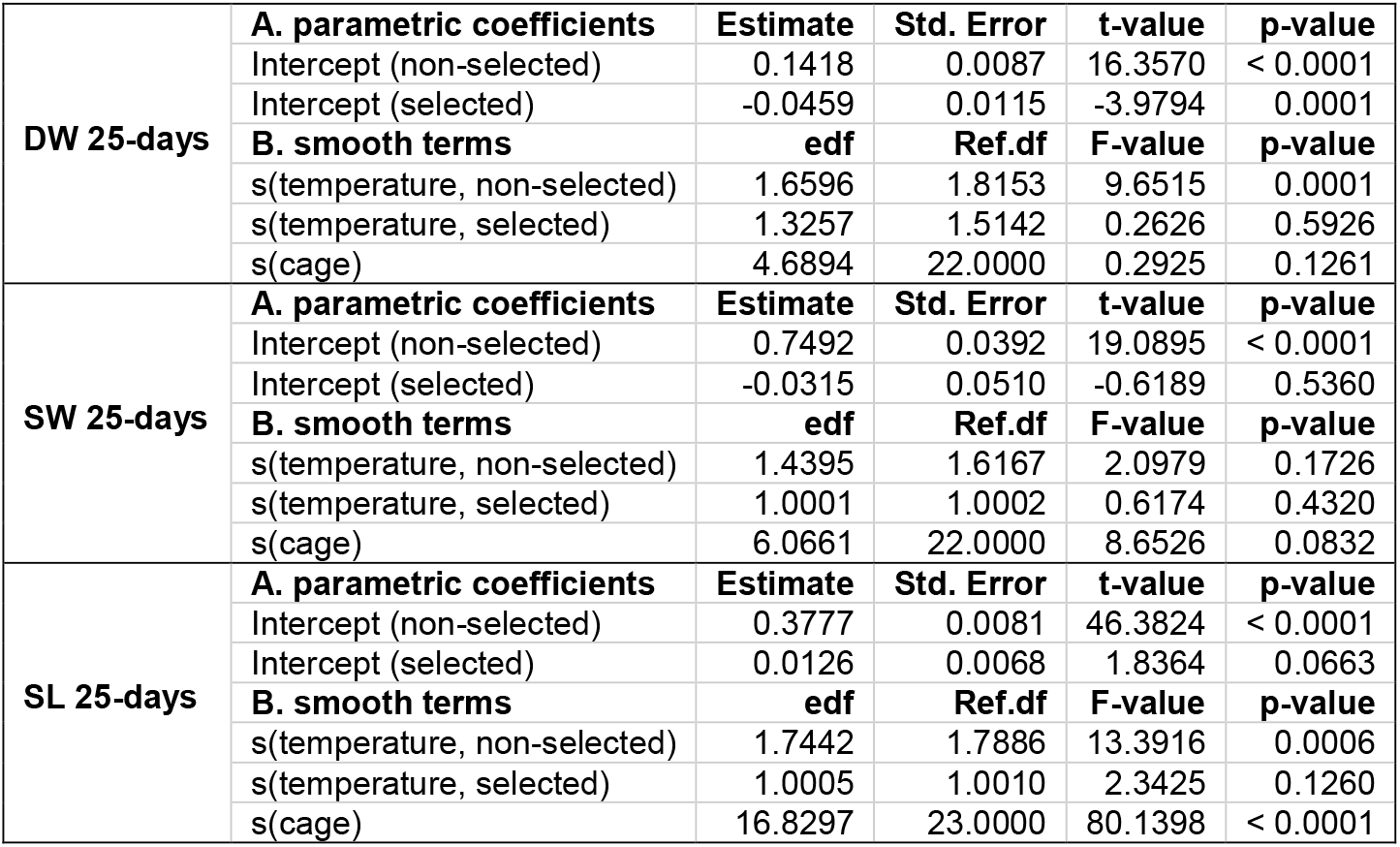
The main and interactive effects of elevated temperature and selection on dry tissue weight (DW), shell weight (SW) or shell length (SL) of *Mytilus* spat following a 25-day challenge. These relations were modelled as additive mixed models assuming scaled t-distributions of residuals. The intercept (or the mean) and the slope or nonlinearity (effective degrees of freedom, edf) of the reference level smoother (naïve) were compared to zero, respectively; and, the treatment level (selected) smoother’s estimates were then compared to the reference level. In addition, the random intercept effects of cage were tested. Significant impacts were considered where p-value ≤ 0.05.

**Table 2:**
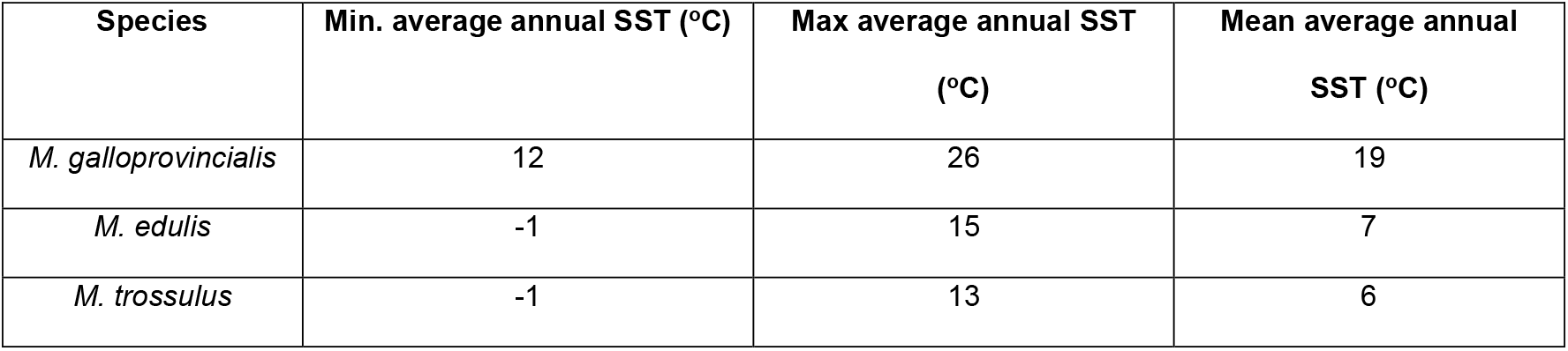
Approximate minimum, maximum and mean average annual SSTs across the geographic distribution of the three *Mytilus* species in the complex: *M. galloprovincialis, M. edulis* and *M. trossulus* from Popovic and Riginos (2020).

**Figure 4:**
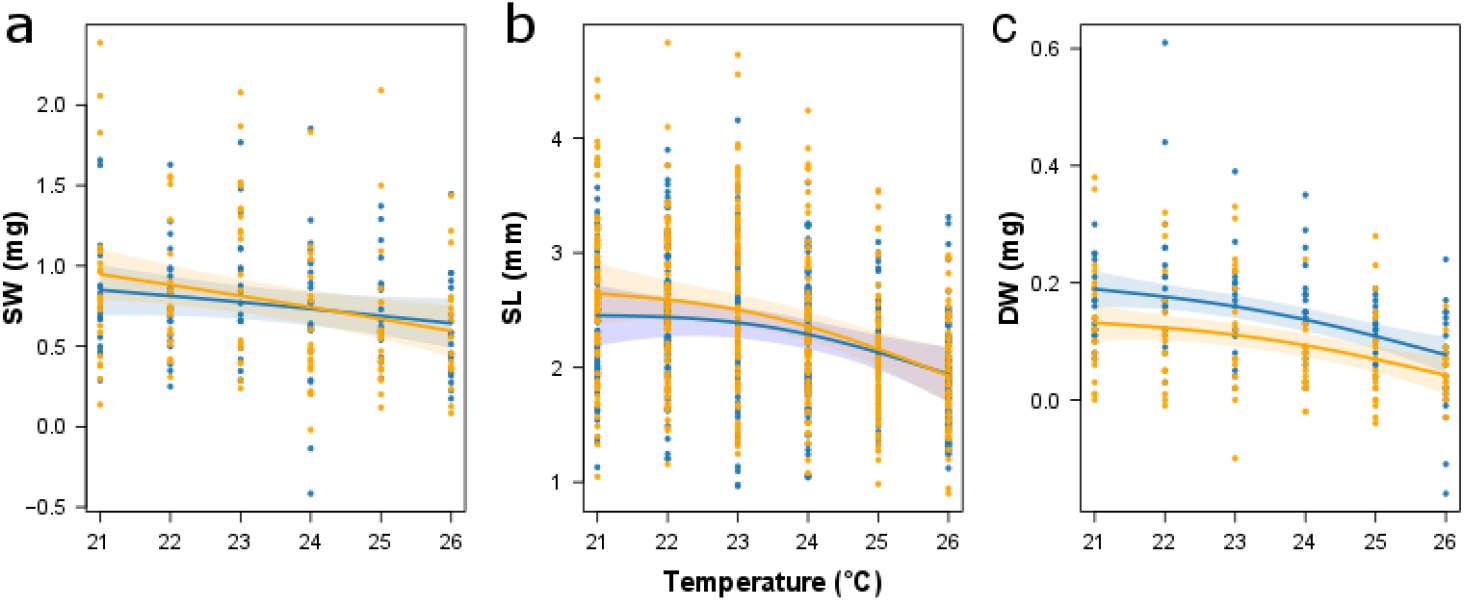
(a) Dry tissue weight (DW), (b) shell weight (SW) and (c) shell length (SL) of selected (S, orange) and non-selected (NS, blue) spat across experimental temperatures (21-26°C), after 25 days of exposure to different temperatures. Each of the data points represents the measurement of an individual spat. These relations were modelled as additive mixed models assuming scaled t-distributions of residuals. Solid lines represent predicted means and shades represent 95% confidence intervals.

## Discussion

In this study, we assessed the impact of strongly elevated summer temperatures on the genetic variation and performance of Baltic juvenile *Mytilus spp*. We found that selection to strongly increased temperatures resulted in an enrichment of *M. edulis* dominated genotypes and concurrently led to reduced tissue growth, in a follow-up growth trial at all tested temperatures. We therefore provide the first evidence that temperature could play an important role in the genomic and phenotypic composition of *Mytilus* mussels in the Baltic Sea. This trend was evident after only a single generation and selection event, and may be indicative of processes that occur in the wild during warm summers. In addition, this rapid response to selection suggests that *Mytilus spp* may be amenable to artificial selection to shift genomic composition and physiological traits of populations, increasing their resilience to ever more extreme global change. More work is needed to understand how genotypes respond to selection in the wild, and to demonstrate if the development of thermally resilient populations via selection in the lab over multiple generations is feasible. However, our results are an important initial step to show that this additional work is necessary and promising.

### Impacts of increased temperature on the genetic variation of mussel spat

We investigated whether exposure to thermal stress can modify the genetic structure of juvenile blue mussel populations by comparing the genetic composition of two spat cohorts, before and after selection, using a multi-species 60K SNP-array (Nascimento-Schulze et al., 2023). The genetic background of the population generally agrees with previously published data on mussels originating from the Baltic Sea, where three species contribute to the population: *M. edulis, M. trossulus* and *M. galloprovincialis* . Nonetheless, the contribution of *M. galloprovincialis* ancestry in spat analysed in this study is considerably higher than has been previously described in this region. While Vendrami et al. (2020) genotyped adult mussels, juveniles (approximately two-months old) were genotyped in our study. The Baltic Sea is a unique environment due to its salinity gradient and limited water mixing with the North Atlantic through the Danish Straight. Individuals possessing *M. galloprovincialis* alleles may be sporadically transported into this region by water currents, carried by ballast water, or attached to other recreational vessels. It is possible that differential survival of genotypes through development could explain the different ancestries. However, more work would be needed to verify this hypothesis.

We find evidence that temperature selection shifted the genetic composition of blue mussels, with higher frequency of *M. edulis* genotypes in the S cohort in comparison to the NS population. Genotype-dependent mortality has been previously observed in mussels (Han et al., 2020; Koehn et al., 1980), supporting the rapid responses to selection in our study. For example, post-settlement selection acting on the *Lap* allele has been observed in *M. edulis* inhabiting a salinity gradient with presence of the allele decreasing through development in low salinities (Koehn et al., 1980). SSTs found across the natural distribution of *M. edulis* and *M. galloprovincialis* are higher than those found within the range of *M. trossulus* (Popovic and Riginos, 2020). These observations support the decreasing proportion of *M. trossulus* alleles in the S population in our study as being driven by selection to temperature tolerant genotypes. Local adaptation to low salinity may have favoured the increase of *M. edulis* genotypes rather than *M. galloprovincialis* in the S population. However, more studies are needed to fully understand the combined effect of multiple stressors, such as temperature and salinity, on selection across *Mytilus* genotypes in the Baltic sea.

On a genomic level, rapid adaptation following a single generation selection event has been previously demonstrated in multiple invertebrate species, including to salinity in larvae of a closely related oyster species, as well as to ocean acidification in mussels (Bitter et al., 2019). Nonetheless, to understand the full impact of this selection on population composition, future work should determine whether this effect persists across time and generations, or whether temporally balanced selection may reduce the impact of this event at a population level (e.g. Durland et al., 2021).

### Impacts of increased temperature on dry tissue weight, shell length and shell mass

Our study found that after 25 days of exposure to elevated temperatures, S juveniles had a lower dry tissue weight compared to NS individuals. In addition, shell length of NS individuals decreased with increasing temperature.

The pervasive impacts of temperature, from molecular to whole-organism levels, affect performance and define the thermal window of a population. Studies on *Mytilus galloprovincialis* show that intermediate warming of 24°C to 26°C, shifts pyruvate kinase to a less active form of the enzyme, leading to hypometabolism and subsequently to metabolic suppression (Anestis et al., 2007). This metabolic suppression enables individuals to lower their energy requirements and allocate surplus energy to the maintenance of basic functions in warm environments (Schulte et al., 2011), and has been observed in Baltic mussels in temperatures beyond the population’s threshold (Vajedsamiei et al., 2021). Therefore, metabolic suppression may have potentially enabled energy to be allocated from growth to physiological processes required to counteract the extreme temperatures. This energetic trade-off may have contributed to the survival of the S cohort during the thermal selection event, but likely at the expense of reduced tissue growth in S individuals.

The higher experimental temperatures, 23°C to 26°C, encapsulate conditions the mussel population will face during the most extreme warm summer periods, from the present day through to the end of the century (Meier et al., 2022; Pansch et al., 2018; Vajedsamiei et al. 2023). Juvenile mussels will unavoidably experience elevated SST and extreme weather events during their development in the Baltic Sea. Our results suggest that these temperatures can exceed the thermal optima for these juvenile mussels. Nonetheless, to confirm whether selection has favoured individuals with lower metabolic rates, oxygen consumption, feeding and excretion rates must be assessed in a follow-up study. Furthermore, to assess whether reduced dry tissue weight demonstrates impaired organism function, the persistence of this phenomenon across to adult organisms must also be investigated.

## Conclusions and Future perspectives

Climate change is projected to modify oceanic physiochemical parameters globally, which have been relatively stable for millennia, with implications for marine ecosystems. Blue mussels are a species group of high-economic and ecological value, thus, guaranteeing that both natural and commercial stocks can thrive under new environmental conditions is a matter of critical importance.

This is the first study to characterise in depth the genetic composition of spat from the Southern coast of the Baltic Sea, using a genomic approach. Using a single generational selection approach to elevated temperature, we observed a clear increase in dominance of *M. edulis* ancestry in the selected population. Our study shows that selection impacts spat performance, with selected individuals able to maintain shell growth at a comparable rate to NS individuals, but at the expense of maintenance of somatic tissue growth. Furthermore, the higher resilience and the fast response to selection of *M. edulis* genotypes to elevated temperature suggest that alleles from this species may confer resilience to increasing temperatures caused by global change, even in highly admixed populations. The differences in the performance and the rapid shift in genetic composition between selected and non-selected cohorts in our study indicate an important role of temperature in this species complex that justifies further investigation. Such information will advance our understanding and ability to quantify the response of blue mussels to thermal stress, as well as their ability to acclimate and/or adapt to such a stressor, in a future ocean. Combining the use of a blue mussel genomic toolbox with knowledge of species-specific physiological mechanisms can ultimately contribute to the development of thermally resilient cohorts, and is a key avenue for further research.

## Data Availability

Raw CEL files are available at https://geome-db.org/query, under the MytiSNP team environment.

Code to run all analysis and produce all figures, and raw admixture output data can be found on Github: https://doi.org/10.5281/zenodo.14362983.

## Acknowledgements

The authors would like to acknowledge the help Ulrike Panknin and Mats Jacobsen in the laboratory. The authors thank Stefano Carboni for the valuable discussions that contributed to the manuscript. The authors acknowledge funding by a NERC GW4+ Doctoral Training Partnership CASE PhD studentship in partnership with CEFAS (awarded to JCNS), funding from the UK Biotechnology and Biological Sciences Research Council (BBSRC) and the UK Natural Environment Research Council (NERC) via a NERC Industrial Innovation Fellowship (NE/R013241/1) and BBSRC Institute Strategic Programme grants (BBS/E/D/30002275 and BBS/E/D/1000207). This work was additionally supported by the Transnational Access program of the EU H2020-INFRAIA project (No. 731065) AQUACOSM -Network of Leading European AQUAtic MesoCOSM Facilities Connecting Mountains to Oceans from the Arctic to the Mediterranean - funded by the European Commission.

