## Supplementary material for "Thermal selection shifts genetic diversity and performance in blue mussel juveniles": Suppl. Table 1

**Supplementary Material (part I)**

**Supplementary table 1:** Measurements of *R. salinas* concentration (cells mL^-1^) over a 72-hour period in 14 L experimental tanks containing no juvenile mussels.

| **Mesocom** | ***R. salinas* 0h** | ***R. salinas* 24h** | ***R. salinas* 36h** | ***R. salinas* 72h** |
| --- | --- | --- | --- | --- |
| M1 | 8768 | 7095.5 | 6912.5 | 6500 |
| M2 | 8021 | 7140.5 | 6825 | 5241 |
| M3 | 7650 | 7417 | 6660.5 | 5504.5 |
| M4 | 7737.5 | 7797 | 7048.5 | 5519 |
| M5 | 8278 | 8257 | 8301 | 7356.5 |
| M6 | 7637.5 | 8477.5 | 7497 | 6488.5 |
| M7 | 9090.5 | 8159 | 7695 | 6138 |
| M8 | 7743 | 7534.5 | 6799.5 | 4943.5 |
| M9 | 7685.5 | 8114.5 | 8798 | 7502.5 |
| M10 | 8278 | 8019.5 | 8689.5 | 6641.5 |
| M11 | 8589 | 9010.5 | 9154 | 7791.5 |
| M12 | 7643 | 7918.5 | 8008.5 | 6659 |

**Supplementary table 2:** Cross validation errors generated by admixture analysis to infer the best number of ancestral populations in dataset containing the selected and non-selected cohorts and the five additional control populations. In this analysis, we tested k’s from a 2 – 10 range. The k with the lower error output is considered the most representative number of ancestral populations for a given dataset.

| **Number of K** | **Cross validation** | **Number of iterations** |
| --- | --- | --- |
| 2 | 0.28594 | 15 |
| **3** | **0.28224** | **20** |
| 4 | 0.28204 | 25 |
| 5 | 0.29493 | 53 |
| 6 | 0.29915 | 34 |
| 7 | 0.30983 | 60 |
| 8 | 0.31806 | 139 |
| 9 | 0.32591 | 64 |
| 10 | 0.34396 | 55 |

**Supplementary table 3:** ANOVA analysis output of impacts of temperature in the shell length (SL) of juvenile mussels either selected (S) or non-selected (NS) to thermal resilience at day-0, prior to the start of the 25-day thermal challenge. In this analysis, a p value lower than 0.05 is considered significant.

|  | **DF** | **Sum Sq** | **Mean Sq** | **F Value** | **Pr (>F)** |
| --- | --- | --- | --- | --- | --- |
| Treatment | 1 | 0.018 | 0.1725 | 0.155 | 0.695 |
| Residuals | 118 | 13.355 | 0.11318 |  |  |

**Supplementary table 4:** Cross validation errors generated by admixture analysis to infer the best number of ancestral populations in dataset containing the selected and non-selected cohorts. In this analysis, we tested k’s from a 2 – 10 range. The k with the lower error output is considered the most representative number of ancestral populations for a given dataset.

| **Number of K** | **Cross validation** | **Number of iterations** |
| --- | --- | --- |
| 2 | 0.34400 | 13 |
| **3** | **0.33842** | **18** |
| 4 | 0.35452 | 37 |
| 5 | 0.37749 | 31 |
| 6 | 0.39126 | 70 |
| 7 | 0.42606 | 51 |
| 8 | 0.44876 | 44 |
| 9 | 0.46886 | 59 |
| 10 | 0.48496 | 29 |

**Supplementary table 5:** Measurements of *R. salinas* concentration (cells mL^-1^) in 14 L experimental tanks over the course of the experiment

| **Tank** | **date t1** | **food t1 (cell ml -1)** | **date t2** | **food t2 (cell ml -1)** |
| --- | --- | --- | --- | --- |
| b23_a | 21.08.2021 | 7770 | 24.08.2021 | 1034 |
| b24_a | 21.08.2021 | 7535 | 24.08.2021 | 2525 |
| b25_a | 21.08.2021 | 6525 | 24.08.2021 | 1716 |
| b22_a | 21.08.2021 | 7615.5 | 24.08.2021 | 3427.5 |
| b24_b | 21.08.2021 | 6977.5 | 24.08.2021 | 4543 |
| b21_a | 21.08.2021 | 6533.5 | 24.08.2021 | 4004.5 |
| b26_a | 21.08.2021 | 6908 | 24.08.2021 | 9201.5 |
| b23_b | 21.08.2021 | 6899 | 24.08.2021 | 1538.5 |
| b21_b | 21.08.2021 | 7148.5 | 24.08.2021 | 3409.5 |
| b26_b | 21.08.2021 | 6807.5 | 24.08.2021 | 3071.5 |
| b22_b | 21.08.2021 | 6561.5 | 24.08.2021 | 3512 |
| b25_b | 21.08.2021 | 6614 | 24.08.2021 | 2358.5 |
| b23_a | 24.08.2021 | 8768 | 27.08.2021 | 887 |
| b24_a | 24.08.2021 | 8021 | 27.08.2021 | 711 |
| b25_a | 24.08.2021 | 7650 | 27.08.2021 | 1432 |
| b22_a | 24.08.2021 | 7737.5 | 27.08.2021 | 912 |
| b24_b | 24.08.2021 | 8278 | 27.08.2021 | 2849.5 |
| b21_a | 24.08.2021 | 7637.5 | 27.08.2021 | 1194 |
| b26_a | 24.08.2021 | 9093.25 | 27.08.2021 | 1690 |
| b23_b | 24.08.2021 | 7743 | 27.08.2021 | 635.5 |
| b21_b | 24.08.2021 | 7685.5 | 27.08.2021 | 1091 |
| b26_b | 24.08.2021 | 8278 | 27.08.2021 | 1307.5 |
| b22_b | 24.08.2021 | 8589 | 27.08.2021 | 710 |
| b25_b | 24.08.2021 | 7643 | 27.08.2021 | 817.5 |
| b23_a | 27.08.2021 | na | 29.08.2021 | 531 |
| b24_a | 27.08.2021 | na | 29.08.2021 | 739 |
| b25_a | 27.08.2021 | na | 29.08.2021 | 1257 |
| b22_a | 27.08.2021 | na | 29.08.2021 | 990.5 |
| b24_b | 27.08.2021 | na | 29.08.2021 | 1453.5 |
| b21_a | 27.08.2021 | na | 29.08.2021 | 1802 |
| b26_a | 27.08.2021 | na | 29.08.2021 | 1213 |
| b23_b | 27.08.2021 | na | 29.08.2021 | 542 |
| b21_b | 27.08.2021 | na | 29.08.2021 | 1857 |
| b26_b | 27.08.2021 | na | 29.08.2021 | 1339.5 |
| b22_b | 27.08.2021 | na | 29.08.2021 | 706.5 |
| b25_b | 27.08.2021 | na | 29.08.2021 | 582 |
| b23_a | 29.08.2021 | 8233.5 | 31.08.2021 | 3211 |
| b24_a | 29.08.2021 | 7962.5 | 31.08.2021 | 2555.5 |
| b25_a | 29.08.2021 | 8136.5 | 31.08.2021 | 4096 |
| b22_a | 29.08.2021 | 7792 | 31.08.2021 | 3318 |
| b24_b | 29.08.2021 | 7647 | 31.08.2021 | 3170.5 |
| b21_a | 29.08.2021 | 7763.5 | 31.08.2021 | 3649 |
| b26_a | 29.08.2021 | 7851.5 | 31.08.2021 | 3167.5 |
| b23_b | 29.08.2021 | 7847.5 | 31.08.2021 | 3207 |
| b21_b | 29.08.2021 | 7265.5 | 31.08.2021 | 3481 |
| b26_b | 29.08.2021 | 8035 | 31.08.2021 | 1653.5 |
| b22_b | 29.08.2021 | 7385 | 31.08.2021 | 2765.5 |
| b25_b | 29.08.2021 | 7633 | 31.08.2021 | 3020 |
| b23_a | 31.08.2021 | 9059.5 | 02.09.2021 | 3081 |
| b24_a | 31.08.2021 | 8962.5 | 02.09.2021 | 3291 |
| b25_a | 31.08.2021 | 8736 | 02.09.2021 | 3006.5 |
| b22_a | 31.08.2021 | 9326.5 | 02.09.2021 | 3354.5 |
| b24_b | 31.08.2021 | 9279 | 02.09.2021 | 4842.5 |
| b21_a | 31.08.2021 | 9278.5 | 02.09.2021 | 4501 |
| b26_a | 31.08.2021 | 9204.5 | 02.09.2021 | 2682.5 |
| b23_b | 31.08.2021 | 9355 | 02.09.2021 | 2856 |
| b21_b | 31.08.2021 | 9318 | 02.09.2021 | 2797 |
| b26_b | 31.08.2021 | 9496.5 | 02.09.2021 | 6323 |
| b22_b | 31.08.2021 | 9160 | 02.09.2021 | 4000.5 |
| b25_b | 31.08.2021 | 9677 | 02.09.2021 | 1492.5 |
| b23_a | 02.09.2021 | na | 04.09.2021 | 3321.5 |
| b24_a | 02.09.2021 | na | 04.09.2021 | 3491 |
| b25_a | 02.09.2021 | na | 04.09.2021 | 3517 |
| b22_a | 02.09.2021 | na | 04.09.2021 | 2363 |
| b24_b | 02.09.2021 | na | 04.09.2021 | 3276.5 |
| b21_a | 02.09.2021 | na | 04.09.2021 | 4019 |
| b26_a | 02.09.2021 | na | 04.09.2021 | 6885.5 |
| b23_b | 02.09.2021 | na | 04.09.2021 | 2030 |
| b21_b | 02.09.2021 | na | 04.09.2021 | 1757 |
| b26_b | 02.09.2021 | na | 04.09.2021 | 6375.5 |
| b22_b | 02.09.2021 | na | 04.09.2021 | 3530.5 |
| b25_b | 02.09.2021 | na | 04.09.2021 | 1289.5 |
| b23_a | 04.09.2021 | 9668 | 06.09.2021 | 4682.5 |
| b24_a | 04.09.2021 | 7655.5 | 06.09.2021 | 2124 |
| b25_a | 04.09.2021 | 8063.5 | 06.09.2021 | 2437.5 |
| b22_a | 04.09.2021 | 8090.5 | 06.09.2021 | 3248.5 |
| b24_b | 04.09.2021 | 7687 | 06.09.2021 | 3030 |
| b21_a | 04.09.2021 | 7961.5 | 06.09.2021 | 4667.5 |
| b26_a | 04.09.2021 | 8066.5 | 06.09.2021 | 8580 |
| b23_b | 04.09.2021 | 7521 | 06.09.2021 | 2148.5 |
| b21_b | 04.09.2021 | 7947.5 | 06.09.2021 | 1169 |
| b26_b | 04.09.2021 | 8222.5 | 06.09.2021 | 6292.5 |
| b22_b | 04.09.2021 | 7943.5 | 06.09.2021 | 1897 |
| b25_b | 04.09.2021 | 8189 | 06.09.2021 | 1133.5 |
| b23_a | 06.09.2021 | 6660.5 | 08.09.2021 | 2368 |
| b24_a | 06.09.2021 | 6262.5 | 08.09.2021 | 1220.5 |
| b25_a | 06.09.2021 | 6714 | 08.09.2021 | 7669 |
| b22_a | 06.09.2021 | 6324 | 08.09.2021 | 4521.5 |
| b24_b | 06.09.2021 | 6351.5 | 08.09.2021 | 2711 |
| b21_a | 06.09.2021 | 6189 | 08.09.2021 | 2813 |
| b26_a | 06.09.2021 | 6724.5 | 08.09.2021 | 6886 |
| b23_b | 06.09.2021 | 7212.5 | 08.09.2021 | 3702.5 |
| b21_b | 06.09.2021 | 4385.5 | 08.09.2021 | 2624.5 |
| b26_b | 06.09.2021 | 6196 | 08.09.2021 | 4074.5 |
| b22_b | 06.09.2021 | 6524 | 08.09.2021 | 2483.5 |
| b25_b | 06.09.2021 | 6102.5 | 08.09.2021 | 2083.5 |
| b23_a | 08.09.2021 | 6491.5 | 09.09.2021 | 1010.5 |
| b24_a | 08.09.2021 | 5946.5 | 09.09.2021 | 874.5 |
| b25_a | 08.09.2021 | 7075.5 | 09.09.2021 | 5939.5 |
| b22_a | 08.09.2021 | 6790 | 09.09.2021 | 4366 |
| b24_b | 08.09.2021 | 6418.5 | 09.09.2021 | 3412 |
| b21_a | 08.09.2021 | 6346 | 09.09.2021 | 1523 |
| b26_a | 08.09.2021 | 7049 | 09.09.2021 | 6547 |
| b23_b | 08.09.2021 | 6707.5 | 09.09.2021 | 1283 |
| b21_b | 08.09.2021 | 6426.5 | 09.09.2021 | 1769 |
| b26_b | 08.09.2021 | 6893 | 09.09.2021 | 3579 |
| b22_b | 08.09.2021 | 6463 | 09.09.2021 | 1935 |
| b25_b | 08.09.2021 | 5936 | 09.09.2021 | 873 |
| b23_a | 09.09.2021 | 8320.5 | 10.09.2021 | 2182 |
| b24_a | 09.09.2021 | 7945 | 10.09.2021 | 1304 |
| b25_a | 09.09.2021 | 8431 | 10.09.2021 | 7490.5 |
| b22_a | 09.09.2021 | 8931.5 | 10.09.2021 | 5316 |
| b24_b | 09.09.2021 | 8630.5 | 10.09.2021 | 3510 |
| b21_a | 09.09.2021 | 8477 | 10.09.2021 | 2061.5 |
| b26_a | 09.09.2021 | 8861.5 | 10.09.2021 | 8661.5 |
| b23_b | 09.09.2021 | 8789 | 10.09.2021 | 1828.5 |
| b21_b | 09.09.2021 | 8124.5 | 10.09.2021 | 2330.5 |
| b26_b | 09.09.2021 | 8478.5 | 10.09.2021 | 4236.5 |
| b22_b | 09.09.2021 | 8424 | 10.09.2021 | 4268.5 |
| b25_b | 09.09.2021 | 7814 | 10.09.2021 | 1839 |
| b23_a | 10.09.2021 | 8519 | 11.09.2021 | 1496 |
| b24_a | 10.09.2021 | 7231 | 11.09.2021 | 1250.5 |
| b25_a | 10.09.2021 | 8521.5 | 11.09.2021 | 6649 |
| b22_a | 10.09.2021 | 9162.5 | 11.09.2021 | 6171 |
| b24_b | 10.09.2021 | 8942 | 11.09.2021 | 2680 |
| b21_a | 10.09.2021 | 8586 | 11.09.2021 | 1803 |
| b26_a | 10.09.2021 | 8141 | 11.09.2021 | 7709 |
| b23_b | 10.09.2021 | 8296 | 11.09.2021 | 1902 |
| b21_b | 10.09.2021 | 9044.5 | 11.09.2021 | 2104.5 |
| b26_b | 10.09.2021 | 9431.5 | 11.09.2021 | 2213.5 |
| b22_b | 10.09.2021 | 8857.5 | 11.09.2021 | 4097.5 |
| b25_b | 10.09.2021 | 7544 | 11.09.2021 | 1421 |
| b23_a | 11.09.2021 | 9301.5 | 12.09.2021 | 1376.5 |
| b24_a | 11.09.2021 | 8272 | 12.09.2021 | 1196 |
| b25_a | 11.09.2021 | 9132 | 12.09.2021 | 7314 |
| b22_a | 11.09.2021 | 9634.5 | 12.09.2021 | 6947.5 |
| b24_b | 11.09.2021 | 8661.5 | 12.09.2021 | 2798.5 |
| b21_a | 11.09.2021 | 9313 | 12.09.2021 | 2162.5 |
| b26_a | 11.09.2021 | 9633 | 12.09.2021 | 9375 |
| b23_b | 11.09.2021 | 9305 | 12.09.2021 | 2211 |
| b21_b | 11.09.2021 | 8739 | 12.09.2021 | 2037 |
| b26_b | 11.09.2021 | 9014 | 12.09.2021 | 2068.5 |
| b22_b | 11.09.2021 | 9337 | 12.09.2021 | 3157.5 |
| b25_b | 11.09.2021 | 9197.5 | 12.09.2021 | 1432 |

**Supplementary Figure 1:** Boxplot of initial shell length (SL, mm) of selected (S, red) and non-selected (NS, blue) spat used in the thermal stress experiment. Each circle represents an individual’s spat SL measurement in millimetres.

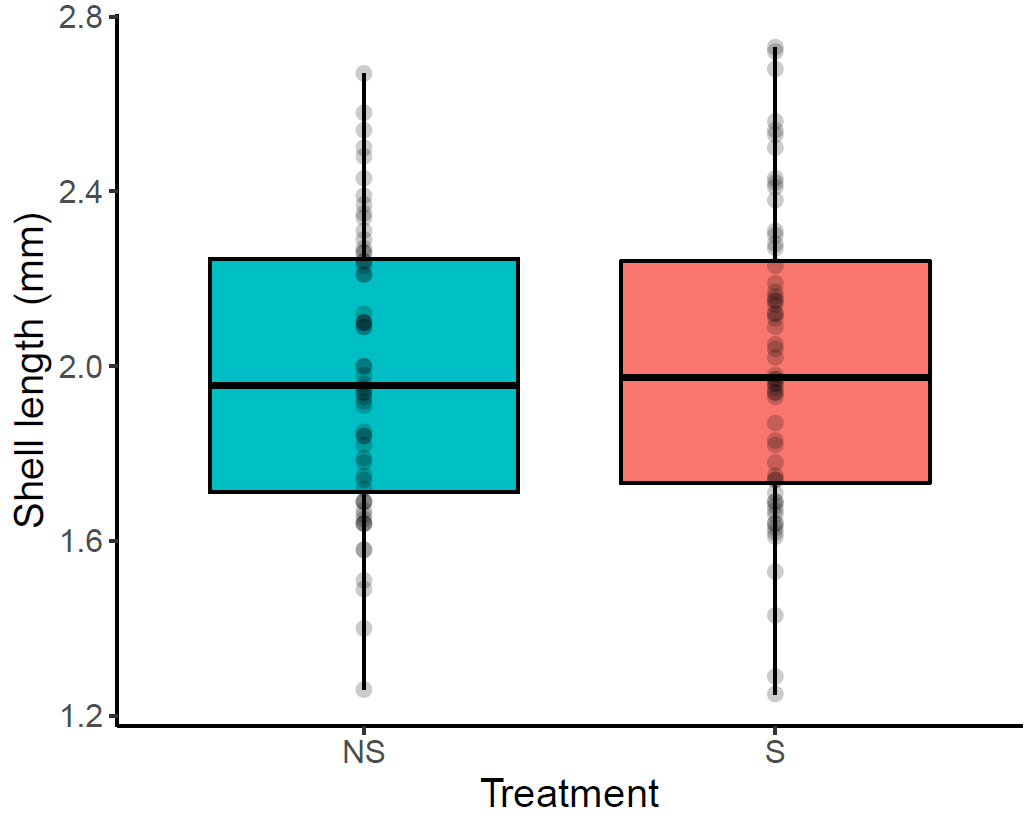

**Supplementary Figure 3:**

Results of genetic admixture analysis using 8,337 SNPs. Each individual is represented by a column partitioned into segments of different colour, the length of which indicate the posterior probability of membership in each cluster. Solid bars represent an individual from a single species background. In this figure we plot a) admixture results of k=3 results for the dataset including both the cohorts (selected and non-selected) in this study. and we the 5 control populations. The dark-blue cluster can be inferred to *M. edulis*, this is the species dominating populations in Budleigh, UK. The light-blue cluster can be inferred to *M. trossulus*, genotypes from this species are highly frequent in finish mussel populations. The yellow cluster can be inferred to *M. galloprovincialis*, as this is the main species colonising Bude, UK.

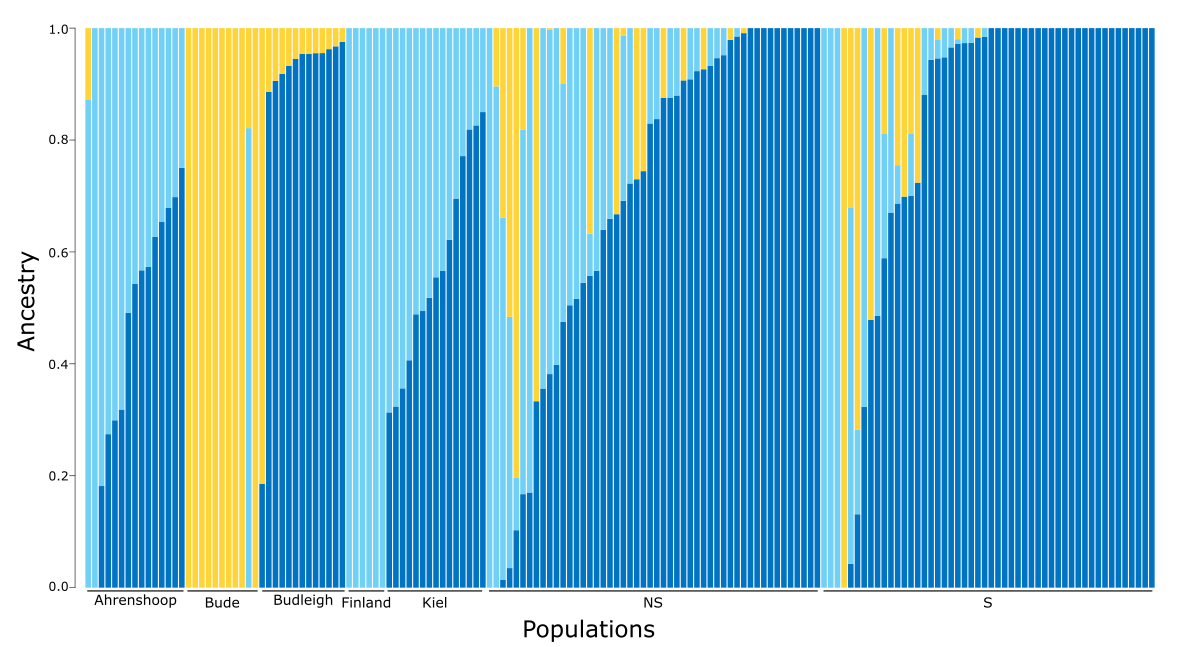

**Supplementary Figure 4:** Results of genetic admixture analysis using 8,337 SNPs. Each individual is represented by a column partitioned into segments of different colour, the length of which indicate the posterior probability of membership in each cluster. Solid bars represent an individual from a single species background. In this figure we plot a) admixture results of a) k=3 and b) k=4 results for the dataset including both the cohorts (selected and non-selected) in this study. and we the 5 control populations. The dark-blue cluster can be inferred to *M. edulis*, this is the species dominating populations in Budleigh, UK. The light-blue cluster can be inferred to *M. trossulus*, genotypes from this species are highly frequent in finish mussel populations. The yellow cluster can also be inferred to M. trossulus sampled 4 years later, as this is the second most frequent species expected in the Selected/Non-selected populations in this study. The red cluster can be inferred to *M. galloprovincialis*, as this is the main species colonising Bude, UK.

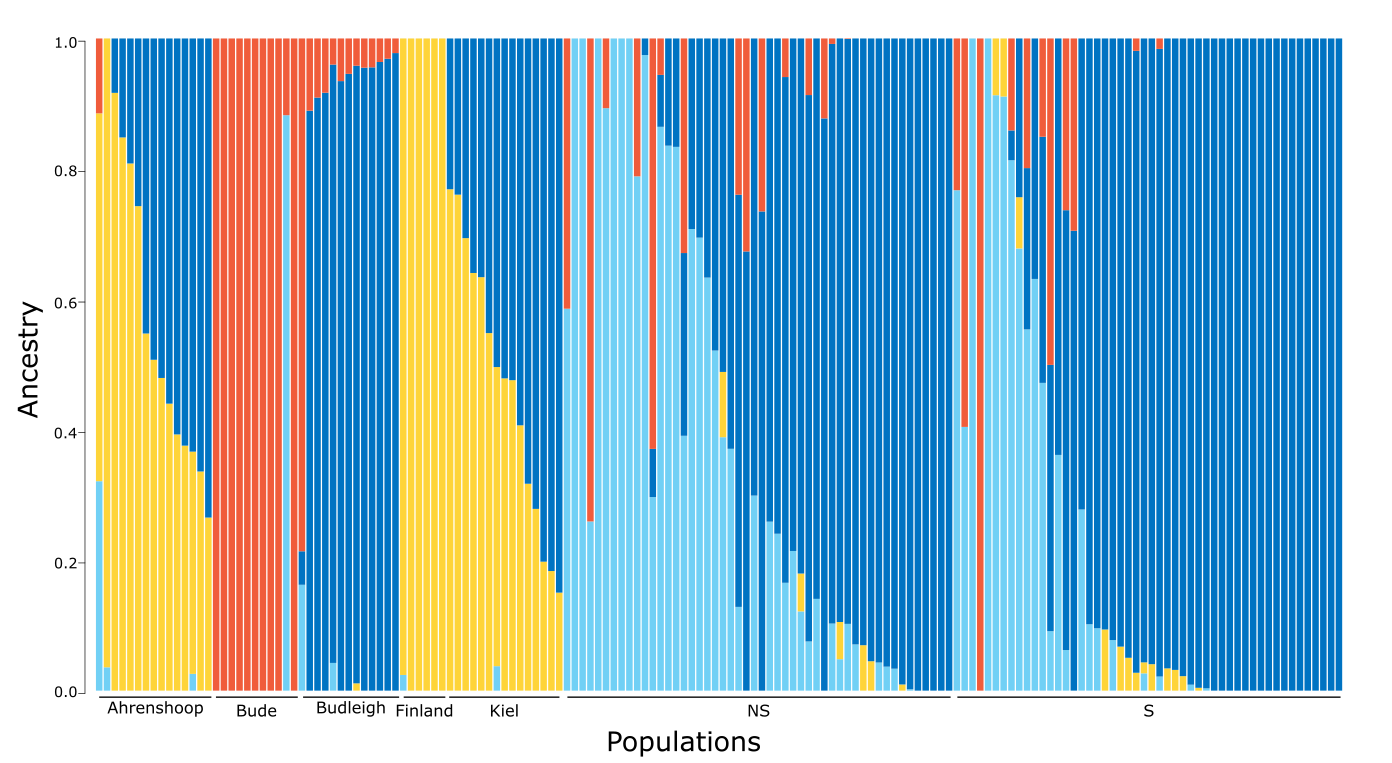

**Supplementary Material (part II)**

Our analysis employed a multi-faceted statistical approach to rigorously investigate the hypothesis that significant differences exist in the ancestry proportions (q1, q2, q3) between the two mussel populations (N vs. NS). This approach included a frequentist Dirichlet regression as well as two Bayesian models, ensuring a robust examination of the genetic compositions across populations.

**Data preparation:** The analysis commenced with the importation of genetic composition dataset containing information on the three ancestry proportions (denoted as q1, q2, and q3) estimated for each sample (N or NS) in the dataset at the best-fitting number of ancestry clusters (K), generated by the admixture analysis. To ensure comparability across samples, the raw ancestry proportions were normalized by dividing each proportion by the sum of proportions for each individual, thereby converting the data into a compositional format suitable for the following analyses. The *population* variable was encoded as a factor.

**Analyses:** A **frequentist Dirichlet regression** approach was adopted initially, allowing for traditional hypothesis testing and p-value interpretation. The *DirichletReg* package facilitated the construction of the Dirichlet regression model, which aimed to capture the differential contribution of ancestry proportions between the two populations. The assumptions underlying the model were evaluated through the examination of standardized and composite residuals, alongside the inspection of residual plots, including histogram and Q-Q plots, to assess the normality and homoscedasticity of residuals.

Subsequently, a Bayesian framework was employed for a more comprehensive understanding of the data's underlying structure, utilizing the *brms* package. Two Bayesian models were specified: one assuming a Dirichlet distribution and the other a logistic-normal distribution for the ancestry proportions. **Bayesian Dirichlet Model** offered interpretability regarding the mean proportions and variability within the compositional data. In essence, the Dirichlet distribution provides parameters that are directly connected to the expected proportions of each part of the composition, facilitating a more intuitive understanding of the data. Then, **Bayesian Logistic-Normal Model** expanded the analysis by allowing for the investigation of correlations between components, making it a robust choice for our analysis aiming to understand the interplay between different ancestry proportions, which could reveal important insights into the genetic makeup and evolutionary processes of the populations under study.

These Bayesian models were fitted with priors tailored to the data characteristics, and Hamiltonian Monte Carlo (HMC) simulations were run to estimate the posterior distributions. To robustly estimate the parameters of these models, we used four chains, each running for 10,000 iterations, which includes a *warm-up* phase of 5,000 iterations to allow for parameter stabilization. The control parameters were carefully chosen to enhance the sampling process: *adapt_delta* was set to 0.99, increasing the step size adaptivity to reduce the risk of divergent transitions, and *max_treedepth* was set to 20, permitting deeper exploration of the posterior landscape. Notably, in these Bayesian models for compositional data, while the primary focus is often on the parts of the composition that show significant differences or effects, the reference component (here *q*1) is also part of the analysis. In this specific model, *q*1's behavior is indirectly inferred because the models estimate the means and variances for *q*2 and *q*3, and *q*1 can be calculated as the remaining proportion, since *q*1+*q*2+*q*3=1 for each observation.

**Bayesian model evaluation and comparison :** Model diagnostics included the inspection of trace plots and density plots for each parameter to ensure proper convergence and the reliability of the posterior estimates. Posterior predictive checks were performed to evaluate the models' predictive accuracy, involving the generation of posterior predictive distributions and their comparison to the observed data through density overlays and statistical summaries.

The relative performance of the Bayesian models was assessed using the Leave-One-Out Cross-Validation (LOO) method, providing a quantitative basis for model selection based on predictive accuracy.

**Results**

**Frequentist analysis with Dirichlet regression**

The frequentist approach utilized Dirichlet regression to assess the differences in ancestry proportions (q1, q2, q3) between the two mussel populations. The results indicated significant effects for the second ancestry proportion (q2) with respect to the population type. Specifically, the estimate for the intercept in q2's model was 0.4825 (std. error = 0.1947, z-value = 2.478 , p-value = 0.0132), suggesting a baseline level for q2 when considering the reference population. Furthermore, the populationS coefficient for q2 was 0.5645 (std. error = 0.2637, z-value = 2.141, p-value = 0.0323), indicating a distinguishable increase in q2's proportion for the S population compared to the reference group. The analysis for q3 showed a positive but not statistically significant association with the population type, with a p-value < 0.0001. The precision model revealed significant variability in the data, suggesting a compositional change across observations.

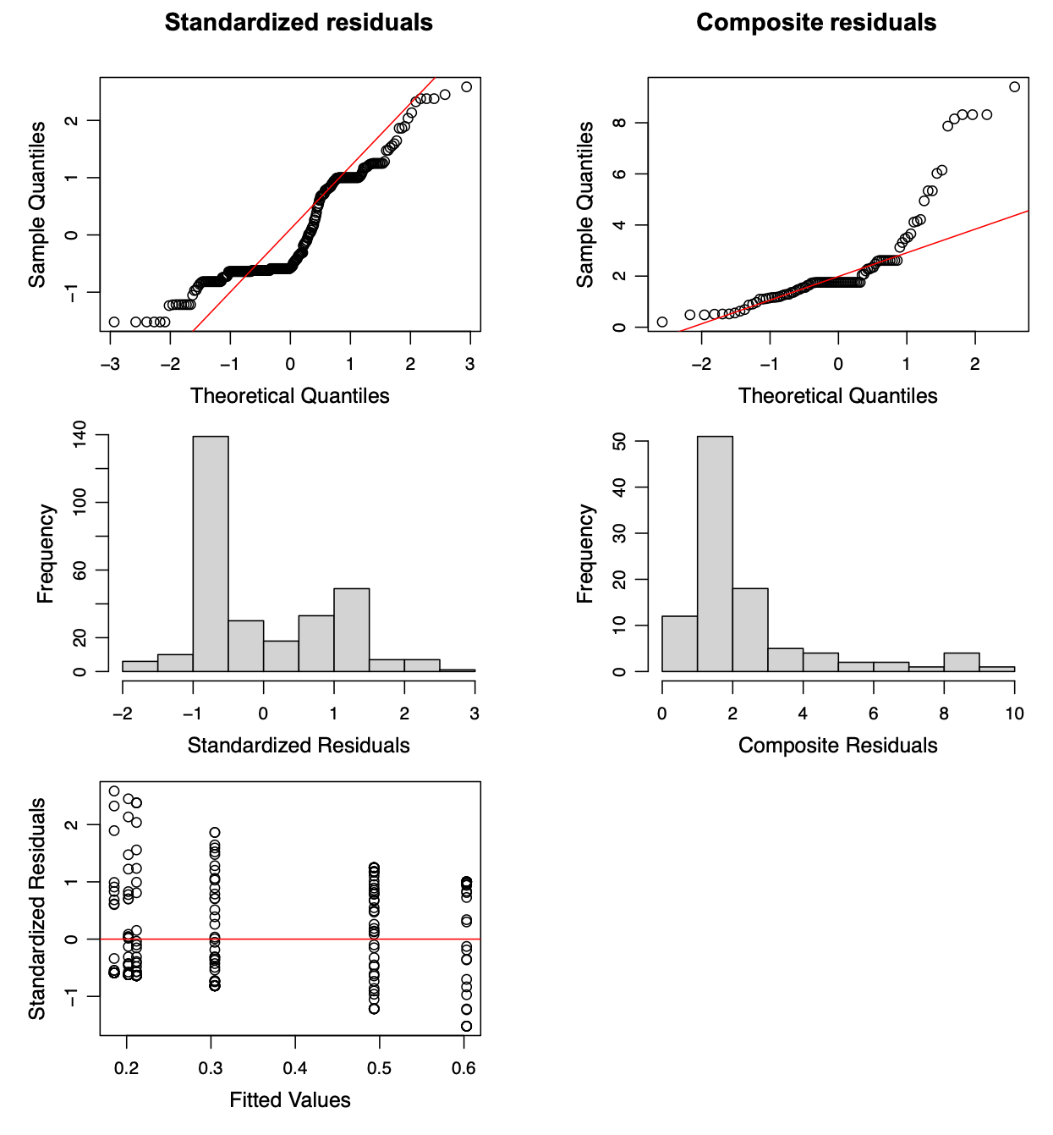

**Supplementary Figure 5:** Evaluation of model fit through residual analysis. The figure illustrates the analysis of standardized and composite residuals to evaluate the fit of the Dirichlet regression model. It highlights the model’s ability to accommodate each compositional component, identifying outliers through standardized residuals, and assessing the overall model fit to multivariate compositional data using composite residuals. Residuals plotted against fitted values and quantile-quantile (Q-Q) plots provide insights into the deviations from normality and homogeneity of variances, crucial for validating model assumptions.

**Bayesian Dirichlet modeling**

For the Dirichlet model with a logit link, the estimated population-level effects indicated that for the second ancestry proportion *q*2, the intercept had an estimate of 0.48 with a 95% credible interval (CI) from 0.09 to 0.87. This suggests a baseline level for *q*2 across the mussel populations. Furthermore, the effect of the **populationS** was estimated to be 0.56 with a 95% CI from 0.05 to 1.08, implying a significant increase in the *q*2 proportion for the **S** population compared to the baseline. For the third ancestry proportion *q*3, the intercept had an estimate of -0.41 with a 95% CI from -0.80 to -0.02, indicating a lower baseline level for *q*3. The **populationS** effect for *q*3 had an estimate of 0.28 with a 95% CI from -0.28 to 0.84, which includes zero, suggesting an inconclusive effect of the population on *q*3.

The 95% CI of [0.44, 0.61] for *ϕ* suggests that we are 95% confident that the true value of *ϕ* lies within this interval. Since the entire interval is greater than 0 but less than 1, it indicates that there is considerable variability within the ancestry proportions, with a tendency towards more extreme distributions. In the context of your study, a �*ϕ* estimate of 0.52 suggests moderate variability within the compositional data. This means that while some individuals might have a balanced mix of ancestry proportions, others might show a dominance of one ancestry over the others.

**Bayesian logistic-normal modeling**

The logistic-normal model estimated the population-level effects for *q*2 and *q*3 as well. For *q*2, the intercept for the reference population (not 'S') was estimated at 2.40 (95% CI: 0.38 to 4.38), and the **populationS** effect was 3.59 (95% CI: 0.74 to 6.42). The estimate indicates a significant increase in the *q*2 proportion for population 'S' compared to the reference population. However, the wide credible interval points to considerable uncertainty about the size of this increase.

For *q*3, the intercept was estimated at -3.20 (95% CI: -5.08 to -1.34), and the **populationS** effect was 1.93 (95% CI: -0.74 to 4.62). The negative intercept suggests that the baseline level for the reference population is low. The positive estimate for population 'S' indicates a higher *q*3 proportion compared to the reference group, but the credible interval includes negative values, implying that the data does not provide strong evidence for an increase in *q*3 due to the population effect.

The family-specific parameters, *σ_q_*_2_ and *σ_q_*_3_, were 7.23 (95% CI: 6.32 to 8.30) and 6.74 (95% CI: 5.88 to 7.77) respectively. This parameter measures the variability in the *q*2 or *q3* proportion across all observations. The high estimate indicates a broad distribution, implying significant heterogeneity across individuals.

The correlation between *q*2 and *q*3 was estimated at 0.49 (95% CI: 0.33 to 0.62). The positive correlation indicates that as the *q*2 proportion increases, there tends to be an increase in the *q*3 proportion as well, within the same population.

**Model convergence and efficacy**

All parameters have *Rhat* of 1.00, suggesting that the model has converged well across all chains. High Effective Sample Size (ESS) values, both as Bulk_ESS and Tail_ESS, for all parameters indicate that the model has effectively sampled from the posterior distribution.

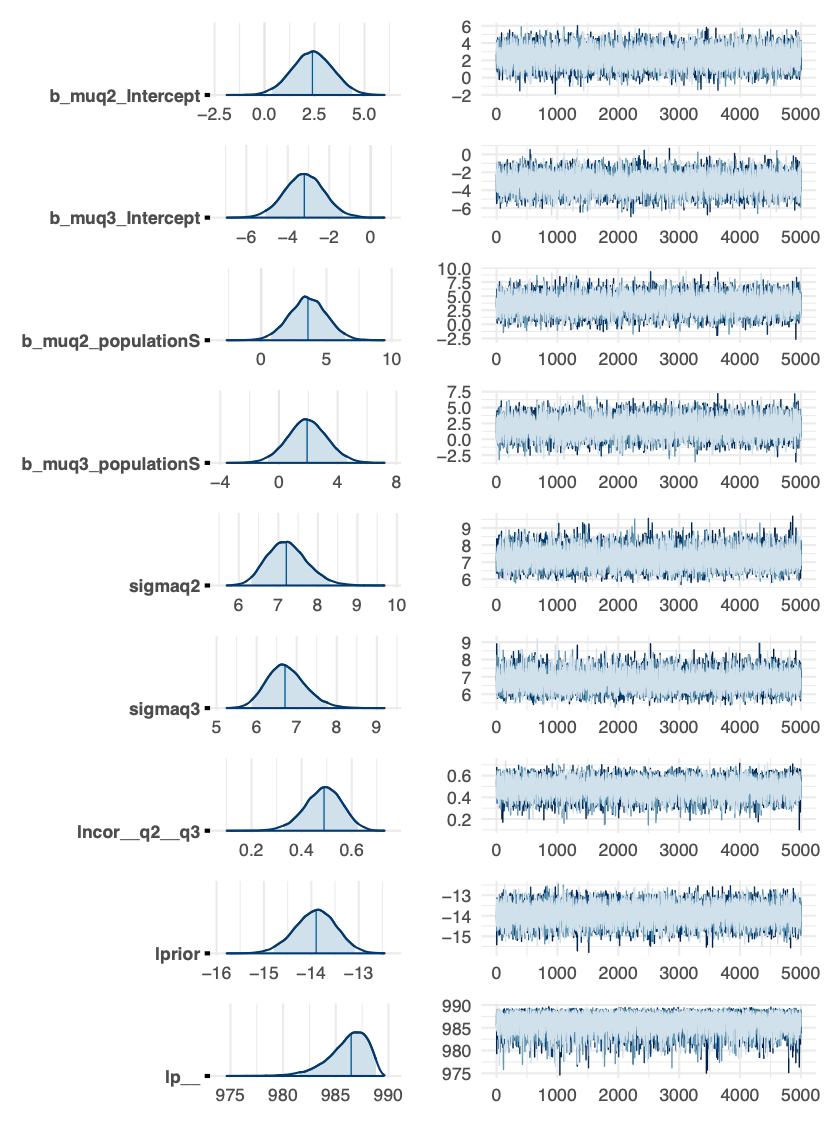

**Supplementary Figure 6:**  Convergence diagnostics for Bayesian logistic-normal model. This figure displays trace and density plots for parameters within the Bayesian Logistic-Normal model, indicating the model’s convergence and the trustworthiness of the posterior estimates. The sampling paths depicted by trace plots confirm effective mixing and equilibrium, essential for accurate inference, while the density plots assess the posterior distributions’ characteristics, ensuring the parameters’ estimates are well-supported by the data.

**Model comparison using LOO**

Model comparison using Leave-One-Out Cross-Validation (LOO) indicated that the logistic-normal model had a higher estimated expected log predictive density (ELPD) with an **elpd_loo** of 992.8 compared to 951.8 for the Dirichlet model. This suggests a better fit of the logistic-normal model to the observed data. The estimated effective number of parameters (**p_loo**) was larger for the logistic-normal model, indicating a more complex model.

Both models showed good performance with all Pareto *k* estimates being satisfactory (all *k*<0.5).

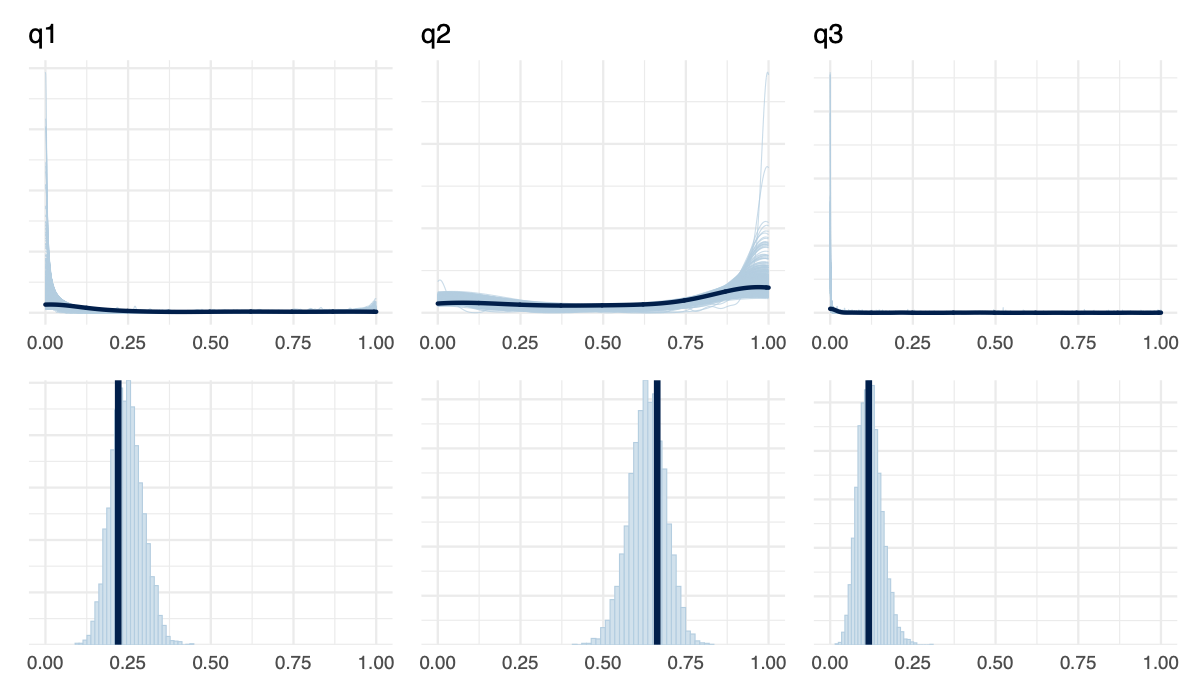

**Supplementary Figure 7:** Posterior predictive checks for the Bayesian logistic-normal model. The figure juxtaposes posterior predictive distributions with actual observations to gauge the Bayesian Logistic-Normal model's predictive precision. Through density overlays (top row), a visual assessment is conducted, complemented by statistical summaries (bottom row) that delve into the model's efficacy, offering a nuanced understanding of its predictive performance and overall reliability.
